# PCR-free, targeted genomic sequencing using Dynamically optimized reference Adaptive Sampling (DORAS)

**DOI:** 10.64898/2026.05.26.727915

**Authors:** Loïc Borcard, Sonja Gempeler, Miguel A Terrazos Miani, Carlo Casanova, Alban Ramette

## Abstract

Whole genome sequencing (WGS) has become a cornerstone of clinical microbiology, enabling comprehensive analysis of microbial genome diversity. However, WGS is often computationally intensive and time-consuming when applied to specific applications like multilocus sequence typing (MLST), where only a subset of genes is only needed for typing. This study evaluates the potential of adaptive sampling (AS), a software-based solution available on Oxford Nanopore Technologies (ONT) devices, to optimize sequencing runs for MLST by reducing the production of unnecessary reads falling outside of the target areas. We demonstrate that AS, when used directly with the target gene sequences, does not reach sufficient target coverage when compared to WGS baseline sequencing due to inefficient read recruitment. Thus, we developed a novel, PCR-free approach, termed Dynamically Optimized Reference Adaptive Sampling (DORAS), which streamlines gene-specific enrichment by targeting genomic regions of interest and their genomic vicinity. DORAS first determines the genomic context of regions of interest for each sample, and then dynamically adjusts the length of the reference sequences during live sequencing. Consensus sequences are periodically constructed and evaluated for taxonomic classification. We demonstrate that full MLST profiles can be obtained in approximately half the time required for whole-genome sequencing to achieve 30X coverage (3 vs. 6 h), with no additional hands-on library preparation time. Validation on clinical isolates from hospital outbreaks belonging to *Corynebacterium diphtheriae*, vancomycin-resistant *Enterococci*, and routine clinical *E. coli* isolates, demonstrated the consistent retrieval of MLST types as compared to standard WGS methods. DORAS thus offers a cost-effective, efficient solution for routine surveillance and outbreak investigations based on MLST types in the clinical setting.

## INTRODUCTION

Targeted sequencing (TS) is an important routine technique in clinical and research settings, offering distinct advantages in terms of clinical utility and cost-effectiveness when compared to Whole Genome Sequencing (WGS) (1). TS panels focus on a particular cluster of genomic regions or a selected number of specific genes. The target genes or regions are typically well known to be related to disease pathogenesis or clinical relevance (2). Several enrichment methods can be applied to increase the concentration of target nucleic acids in a sample via PCR on probe-based enrichment (3) or CRISPR-based strategies (4, 5) making them more amenable to Next Generation Sequencing (NGS). Beside PCR-based, probe-based and CRISPR-based TS, which are all biochemical procedures, Oxford Nanopore Technologies (ONT)-based Adaptive Sampling (“AS” or “Read Until”) is a computational technique that rejects non-target reads during live sequencing, ensuring only relevant reads are fully sequenced (6, 7). While ONT’s built-in AS implementation provides a convenient option for enriching or depleting large, predefined target regions e.g. (8), its current lack of customization, specifically the inability to adjust targets or set depth limits during a run, limits its application to specific, predetermined scenarios. Other AS software implementations which are not embedded directly in ONT devices, such as readfish (7, 9) or BOSS-RUNS (10), can offer more flexible customization, but may encounter compatibility issues with newer versions of the ONT sequencing devices and software (2).

In the context of bacterial outbreak investigations, robust microbial typing is essential for tracking pathogen spread and understanding genetic relationships between pathogenic isolates. Multilocus Sequence Typing (MLST) remains a cornerstone of outbreak surveillance because of its reliability and ability to differentiate closely related strains (11– 13). The use of seven to ten housekeeping genes for sequence typing (ST) allows MLST to provide strong discriminatory power, while delivering results that are unambiguous and easily comparable (11–13). Although the latter does not provide the highest discriminatory power as compared to core genome MLST or SNP-based analyses (14, 15), MLST-based typing is still used to classify isolates below the species levels for outbreak confirmation, surveillance or for understanding transmission dynamics, e.g. (16, 17).

Currently, identifying the small (<500 bases) genomic regions of interest (ROI) required for ST identification relies on two primary methods, each with distinct limitations: The standard approach involves a PCR step with species-specific primers typically followed by Sanger sequencing of the generated amplicons. This method relies heavily on primer quality and requires separate sequencing for each ROI, which becomes costly at scale e.g. (18). The modern alternative is WGS, where the entire genome is sequenced at sufficient depth to extract the necessary ROI. While WGS provides an exhaustive protocol that overcomes the low representation of reads often found in small ROIs, the expensive approach may generate unnecessary sequence data, particularly for targeted applications like MLST screening, where non-ROI sequences are not directly exploited. The efficiency and associated cost of the chosen molecular approach becomes even more paramount during outbreaks involving numerous clinical isolates. Public health officials and clinicians require tools that allow them to quickly identify clusters and track bacterial spread without compromising on accuracy or comprehensiveness. Consequently, there is a pressing need for a fast, cost-effective approach to sequence small ROIs that avoids the labor intensity of Sanger sequencing or the inefficient sequencing of entire genomes.

Here, we developed a method that provides MLST profiles while maintaining flexibility for varying batch sizes and eliminating the need for PCR. To this end, we developed Dynamically Optimized Reference with Adaptive Sampling (DORAS), a workflow that leverages the ONT live data stream and tailors the data generation process specifically for MLST. We first assessed standard AS performance against WGS baseline for MLST data production. We then established the novel PCR-free DORAS approach on a model organism (*E. coli*) before testing it on common outbreak species, including *C. diphtheriae* (CDIP), vancomycin-resistant *E. faecium* (VRE), and routine clinical *E. coli* isolates. Our results suggest that DORAS speeds up the turnaround time (TAT) by two to five times and reduces per-sample costs by three times compared to Illumina WGS and two times compared to standard Nanopore-based WGS.

## METHODS

### Sample origin, cultivation, DNA extraction

All isolates were collected by the clinical microbiology laboratory of the Institute for Infectious Diseases (IFIK), University of Bern, Switzerland. The clinical samples of *E. coli* were obtained from anonymized positive blood cultures in 2025 from routine diagnostics. *Escherichia coli* strain ATCC 25922 was used as a reference. All CDIP isolates were associated with local outbreaks in Switzerland from July to September 2022, and were described in a previous study (17). The VRE isolates were associated with local hospital outbreaks in a Swiss hospital, and were collected between 2017 and 2021. They were isolated on CHROMagar VRE plates (CHROMagar, Paris, France) during routine analysis. All isolates in the study have been anonymized and no patient information is used in the interpretation of the bacterial genomic data in this study. Publication of this analysis does not harm or influence neither patients nor institutions. Ethical committee approval was therefore not requested. All isolates were stored at −80°C, and re-grown on CSBA plates before DNA extraction. Genomic DNA was extracted from agarose plates using PureLink Genomic DNA kit (Thermofisher, Switzerland), or with Maxwell RSC Cultured Cells DNA Kit (Promega, Switzerland). The WGS and DORAS analyses were done on the same genomic DNA extracts.

### Library preparation and GridION sequencing for genomic DNA

When indicated, samples were sheared using g-Tubes (Covaris, UK) at 21,000 g. Otherwise, native gDNA was used following genomic extraction. Nanopore libraries were produced with the rapid barcoding chemistry SQK-RBK114.96 (RBK), according to the manufacture’s recommendations. The libraries were loaded onto standard FLO-MIN114 flowcells and sequenced in batches of 20 samples for 2-72 h on GridION X5 sequencer with real-time basecalling and under high accuracy mode (HAC). All experiments were performed on MinKNOW (v24.11.8) and using Guppy (v7.6.7+de8939544).

### DORAS workflow

The DORAS pipeline consists of two sequential phases (**Fig. 1**): **Phase 1 – Reference-extension (using live-produced data)**. The initial step of the “extension-phase” establishes the optimal extension length for each sample. To do so, several simulations of the model are run (see Methods section) to determine the optimal quantile (and reference extension) to be used based on the distribution of the read length. Once the optimal extension length is obtained, the actual extension process can begin. Reads are concatenated and mapped to the template reference containing the target *loci, e*.*g*. seven for the MLST scheme of *E. coli*. For each target gene, the reads enabling a genomic extension beyond the 5’ and 3’ ends of ROI are identified (**Fig. 1**). As a starting point, the generated reads are mapped to the initial reference provided (species specific MLST loci). Reads that map but contain an overhang (distance between the last mapping position and the end of the target read) that is greater than 0.8 times the mapping region are discarded (19). After filtering, the longest extension is selected, and 500 bp are trimmed from each end (default value). Among the remaining reads, the read providing the longest extension on either 3’ or 5’ end is identified and used to extend the previous sequence (**Fig. 1**). During the extension phase, the initial reference provided is mapped to the current extended reference. The start position of the mapping corresponds to the current extension length at the 5’ end (upstream), and the difference between the end of mapping region and the length of the current reference is considered as the 3’ (downstream) extension. The final FASTA file of each sample is concatenated into a single file containing all final references. Because the extension relies exclusively on reads generated in real time, the workflow can dynamically expand the genomic context of each ROI without any prior knowledge of the surrounding sequence, thereby producing an “Optimized Reference” that is tailored to each isolate.

**Figure 1.**
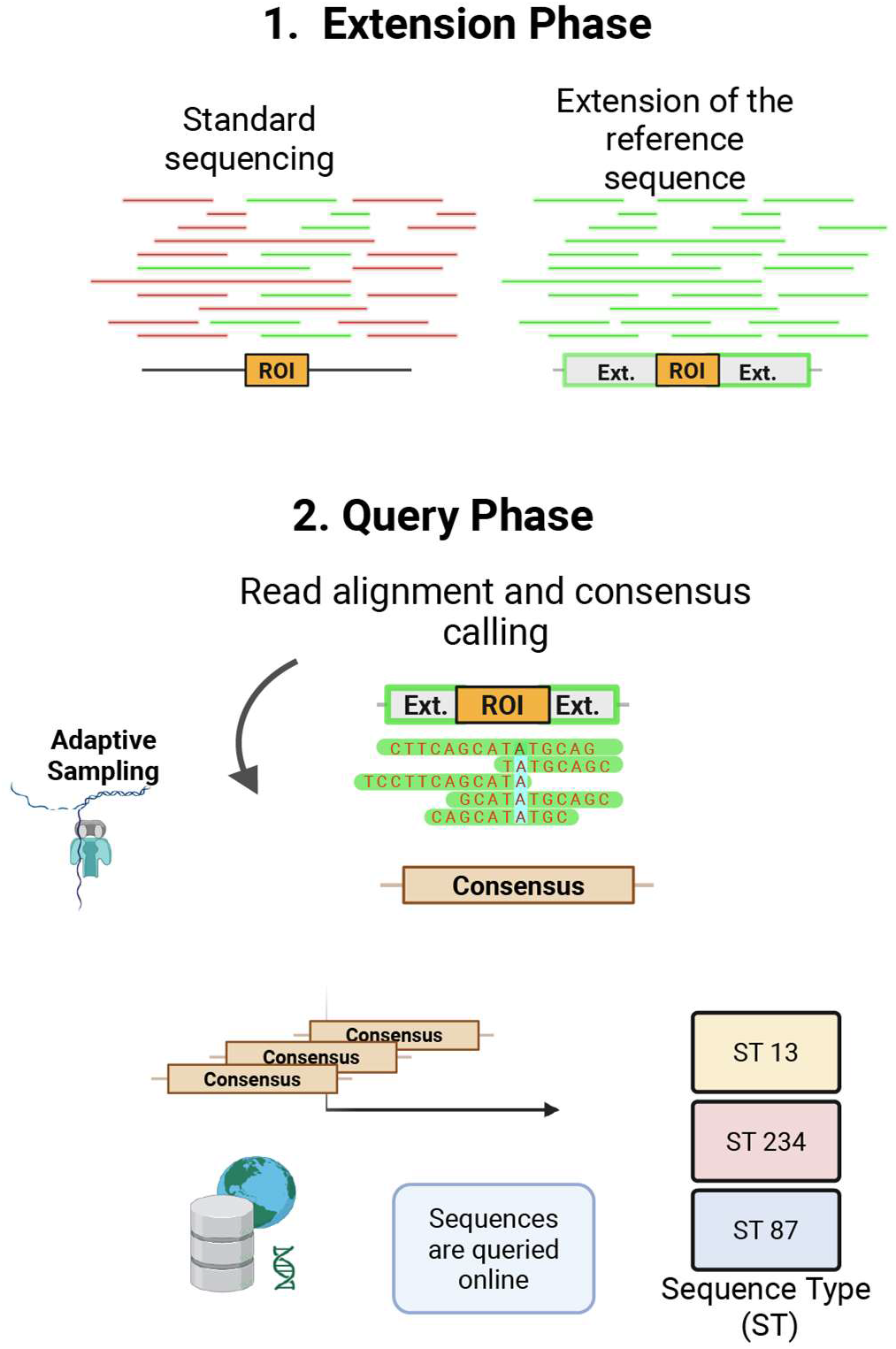
Overview of the DORAS protocol. The data livestream from the ONT device is used to extend the initial reference (here MLST *loci*) (Phase 1: Extension phase). At the end of phase 1, the optimized reference allows the enrichment of the *loci* to quicken the consensus calling process (Phase 2: Query Phase). The process is terminated once all ST have been resolved. Consensus calling only begins once sequencing mean coverage reaches 20X.

#### Phase 2 – Query (AS enrichment; Fig. 1)

The optimized reference FASTA is supplied to the ONT device for adaptive sampling. The accumulating reads are periodically remapped to generate consensus sequences with Medaka (v2.1). After each consensus building round, the depth of each ROI is verified and once over 20 for all samples, the FASTA is sent to the BIGSdb web API (11, 20) for sequence-type (ST) identification. The query phase continues until all consensus sequences reach a minimum depth of 20 upon which the consensus sequences will be queried with the appropriate BIGSdb and the process stop once all STs are resolved (all *loci* match perfectly).

### Analysis of coverage over time

Coverage analysis was performed by processing the raw sequencing data. First, all generated reads (FASTQ format) were collected. For AS runs, we specifically extracted the subset of reads that were marked as “accepted” by the AS software, using the classification provided in the sequencing summary and specific AS output logs. For the control WGS runs, all valid reads were retained for analysis without this filtering step. The reads were then mapped using minimap2 (v2.11) to the species-specific MLST loci consisting of *adk, icd, mdh, purA, gyrB, fumC*, and *recA* for *E. coli, atpA, adk, purK, pstS, gdh, gyd*, and *ddl* for VREs, and *leuA, dnaE, atpA, dnaK, rpoB, fusA*, and *odhA* for CDIP. Mapping information was extracted and converted into a tabular format using seqkit (v2.6). Those tables were then merged with timestamp data extracted from the original FASTQ headers, also using seqkit, to associate each alignment with its specific generation time during the run.

Data analysis and visualization were performed using the R programming language (v4.2), utilizing tidyverse (v2.0.0), data.table (v1.16.4), and ggpubr (v0.6.0) packages. The alignment length for each aligned read was used to calculate the cumulative mean coverage for each barcode over the duration of the sequencing run. This longitudinal coverage data was then used to determine the specific time points at which each sample reached milestone mean coverage values from 10X to 40X.

### Mathematical modeling of AS dynamics

The capacity of the reference provided to Read until (the API powering AS) to recruit reads containing the ROI depends on read length distribution. The latter is typically right skewed, and modelled by a Gamma distribution (21, 22). In our case, we aimed to enrich for target ROI whose size is below the quantile 99 of the read length distribution (N50 of 10kb) and thus can be fully covered by a single read. Because the ROI may lie within a read but far away from its ends, such cases are less likely to be detected and sequenced appropriately. While the concept of “buffer size” is mentioned in the ONT technical documentation (23), detail about the extension size (“buffer size”) is not provided for small ROI and may highly depend on the sample conditions. In fact, an overly long extension around the ROI can conceptually be detrimental to the enrichment process by recruiting reads that do not contain the ROI, thereby reducing the efficiency of the enrichment (see Results section).

In an effort to understand the complex relationship between these parameters and determine the optimal reference extension length that maximizes target coverage, we developed a stochastic model simulating the AS process on ONT devices (full mathematical model specification provided in **Supplementary Material**). The model treats the flow cell as a collection of independent pores (default 500). Read length (*L*) is extracted from a FASTQ file or generated from a gamma distribution. The model simulates the AS decision-making process at each pore and fixed number of ROI (*c*_ROI_). The initial step (1) computes, given a read length distribution, the extension size based on a chosen quantile (e.g., 0.99). (2) Using the newly computed reference length the set of all possible combinations of subsequent (k-mers, *K*(*L*)) of length (*L*) from the entire genome (*K*_*total*_(*L*)), extended reference (*K*_ref_(*L*)) and the ROI. The ratio (*r*_*ref*/*total*_ = *K*_ref_(*L*)/ *K*_*total*_(*L*), represents the frequency of k-mers mapping to the extended reference which enables the (3) prediction the number of reads mapping to both the extended reference after a specific time. These k-mers correspond to the number of reads enriched during the AS process. The final count of k-mers that map to the ROI corresponds to a subset of k-mers that map to extended reference. The number of k-mers mapping to the ROI is estimated using a the computed *K*_*roi*_/*K*_*ref*_ which is the frequency of the k-mers mapping to both the ROI and the extended reference.

### Estimation of ideal reference length for each sample

Each clinical sample has a specific read length distribution and thus the reference length must be adapted. Using our model, we simulate for each sample a 30-min run with read length distribution as input to obtain a yield of reads given a chosen quantile. We simulate several runs with a range of ten different quantiles from 0.5 to 0.99. The quantile reporting the highest number of reads is chosen for this specific sample.

### Data and code availability

Raw sequencing data of this study have been deposited in BioProject PRJNA1440396. Individual accession number for each sample is provided as Supplementary Tables S1-S4.

DORAS is implemented in Python and is available at https://github.com/RametteLab/DORAS. We provide a pixi environment including the necessary Python dependencies: numpy (v1.26), medaka (v2.1.1), mappy (v2.24), samtools (v1.21), pandas (v2.3.3), pysam (v0.23.0). A copy of the version at the time of publication has also been deposited at Zenodo https://doi.org/10.5281/zenodo.20391868.

### Usage of large language models

During manuscript preparation, Microsoft Copilot was used to assist with language editing and refinement of selected passages. Qwen3-Coder-Next(24), accessed through Ollama (25), was used to assist with selected code-development tasks. The authors reviewed, edited, and verified all AI-assisted outputs and take full responsibility for the final content of the manuscript and code.

## RESULTS

### Default AS produces low coverage of target sequences

We first conducted a baseline experiment using *E. coli* ATCC 25922 (accession number: NC_000913.3) in a multiplexed experiment using four barcodes, as technical replicates. In the AS run, we supplied the MinKNOW software with the sequence of the corresponding seven Achtman MLST *loci* (26). We compared total read output and ROI coverage after the first two hours of sequencing between WGS and standard AS (**Fig. 2A**). Under WGS, 235 k reads were produced after 2 h, of which only 1,318 mapped to the seven ROI across all samples (**Fig. 2B; Table 1; Table S1**). The grand mean coverage for all four barcodes was 48X under WGS conditions. The standard AS run generated 5.2 M reads, yet only 714 (0.11 %) aligned to the MLST ROI, yielding an average per-locus coverage of 12X across the four barcodes. Under standard WGS conditions, 0.56% reads mapped on average to the seven ROI, while the percentage was only of 0.11% with AS (**Table 1**). The percentage of mapping reads under WGS conditions represents the expected default levels of reads containing the ROI. Thus, the ratio of 5.1 (*i*.*e*. 0.56/0.11) between WGS and standard AS ROI mapping percentages suggests that the default AS was on average 5 times less efficient at recruiting reads mapping to the ROI as compared to WGS (**Table 1**). This emphasizes that using the short ROI sequences as a reference does not allow an efficient recruitment of reads containing the target genes if they are located beyond the first 500 bases of either end. This thus prevents the enrichment of molecules that may also contain the ROI (**Fig. 2A**). To circumvent this issue, we thus suggest extending the target region upstream and downstream of the ROI to detect reads that potentially contain the ROI.

**Table 1.** ROI mapping statistics using WGS and default AS (*E. coli* ATCC 25922).

| Sample | WGS |  | AS |  | Fold difference<br>in mapping to<br>the ROI<sup>3</sup> |
| --- | --- | --- | --- | --- | --- |
|  | Total<sup>1</sup> | Mapping to<br>ROI<sup>2</sup> (%) | Total | Mapping to ROI<br>(%) |  |
| Barcode01 | 53,814 | 380 (0.71%) | 1,285,936 | 1322 (0.10%) | 7.1 |
| Barcode02 | 60,506 | 485 (0.80%) | 1,549,230 | 1792 (0.12%) | 6.7 |
| Barcode03 | 59,774 | 206 (0.34%) | 1,061,963 | 979 (0.09%) | 3.8 |
| Barcode04 | 61,087 | 247 (0.40%) | 1,285,423 | 1461 (0.11%) | 3.6 |
| Total | 235,181 | 1318 (0.56%) | 5,182,552 | 5554 (0.11%) | 5.1 (sd 1.9) |

<sup>1</sup>Total number of reads per barcode. <sup>2</sup>Number and percentage of reads mapping to the ROI. <sup>3</sup>Fold difference in percentage read mapping to the ROI between WGS and AS (WGS used as reference).

**Figure 2.**
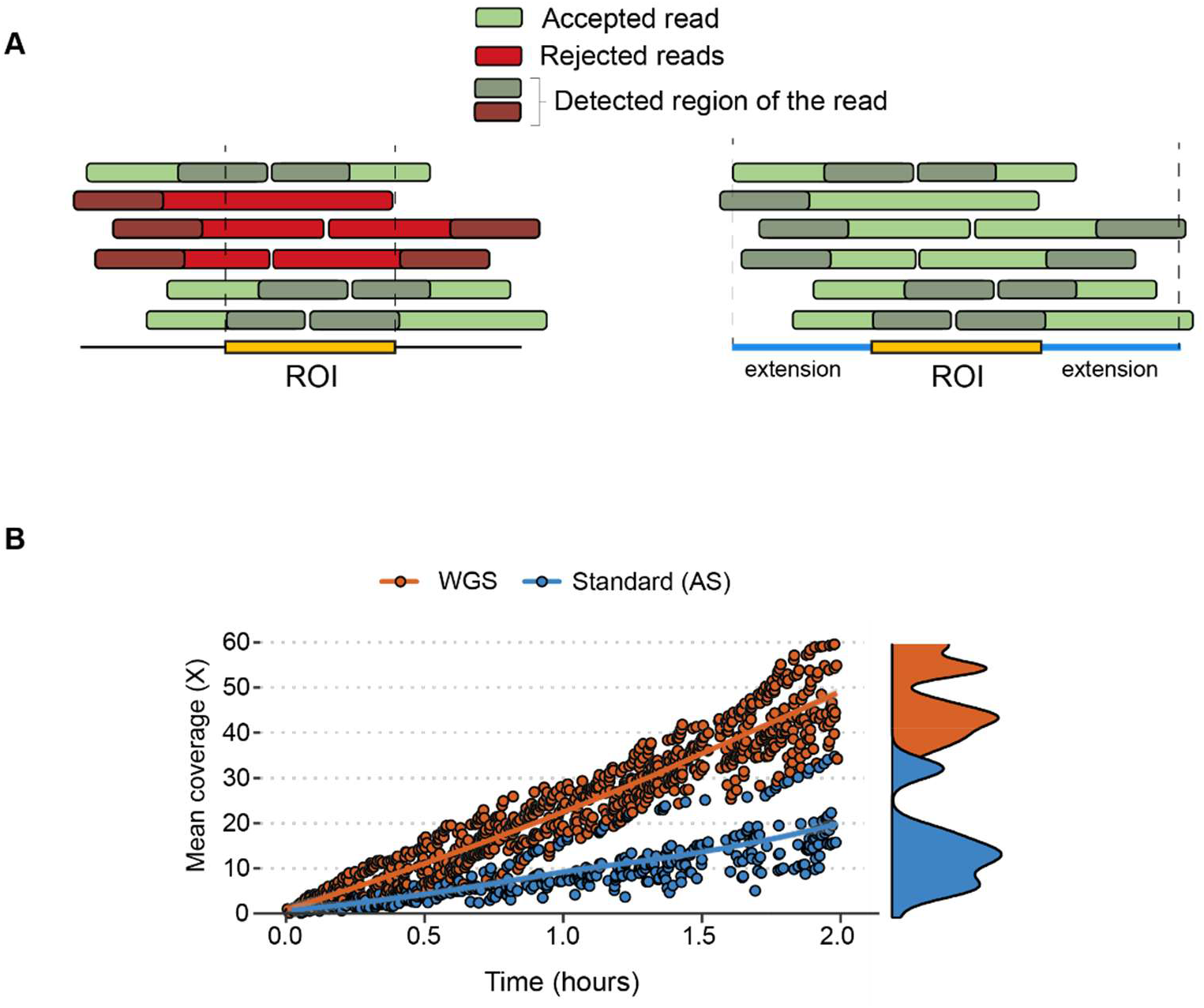
Optimizing AS selection parameters to maximize target gene coverage. **A**) Schematic overview of the AS selection process. Based on the provided ROI sequence, only the first 500 bases translocating into a pore are mapped (“detected region”, greyed-out in the drawing). This implies that only reads containing the ROI at one end will be detected (green) and the other reads will be rejected (red) even if they contain the ROI. **B**) The reference sequences (ROI) of each gene are extended upstream and downstream by up to 25 kb from the ROI. The extension of the reference enables the capture of reads whose detected regions did not initially map to the ROI. **C**) Comparison of the mean coverage for the *E. coli* specific loci obtained for the first two hours of sequencing with WGS and standard AS.

### With live-generated data, the DORAS workflow can dynamically expand the genomic context of the ROI

Increasing the genomic context around the ROI is necessary to avoid falsely rejecting reads that contain the ROI but whose detected region falls outside the ROI (**Fig. 2A**). Extending the reference beyond the ROI must be sufficient to capture reads of all lengths that contain the ROI at any position (**Fig. 2B**). However, an over-extension of the reference sequence can counter-balance the benefits of AS, as more off-target reads may lead to longer sequencing time and not necessarily to additional information for the resolution of the correct ROI sequences (**Fig. S1**). Each of these additional reads not mapping to the ROI are occupying the pores, thereby preventing other molecules from being sequenced. To avoid over-extension, it is necessary to accept only up to the longest reads that can contain the ROI. A robust strategy could consist of computing the 99th quantile of the read length distribution (**Fig. 1**), and extending the reference sequences on both sides accordingly (referred to as “Optimized Reference” from this point forward). The 99th quantile of read length captures most reads containing the ROI, but if the distribution is long-tailed, this value can become excessively large, resulting in sub-optimal enrichment conditions, as detailed in the next section.

### Modeling of optimal reference sequence size

To understand the relationship between coverage of the ROI with read length and sequence length, we built a model of AS that accounts for the distribution of read lengths (**Fig. S2**), modeled here by a gamma distribution (21, 22). This model allows us to estimate the optimal extension size. In the current case, we consider seven ROIs, each 500 bases long. The model simulates the probabilistic nature of AS. As output, the simulation displays the mean of the distribution of the reads mapping to the ROI (here three simulations for each condition are shown). Using a grid search that iterates over a range of read lengths (2-20 kb), one can determine the optimal quantile that generates the maximum mean count for a given read size (**Fig. 3A**): For instance, the read size of the DNA sample of ATCC *E. coli* after Covaris g-TUBE shearing (see Methods section) corresponded to ~2.4 kb on average and the same sample without shearing produced 4.9-kb read length on average, the optimal quantile being 0.99 and 0.90, respectively. Our model focuses on the impact of the length and aims to simulate the effect of the reference length given a certain read length distribution (**Fig. 3B**): To compare the simulation with experimental data we used the length of reads under the aforementioned conditions as input for the model to estimate the read count (mapping to the ROI). The comparison was limited to the first hour of the run to reduce the influence of the pore count decreasing over time under each condition. While our model underestimated read counts under certain conditions this inconsistency is not highly relevant in our case since we want to evaluate fold change differences between the different conditions. This model could be used during a sequencing run with the generated reads to determine the optimal condition under which AS can enrich ROI for each specific sample for instance.

**Figure 3.**
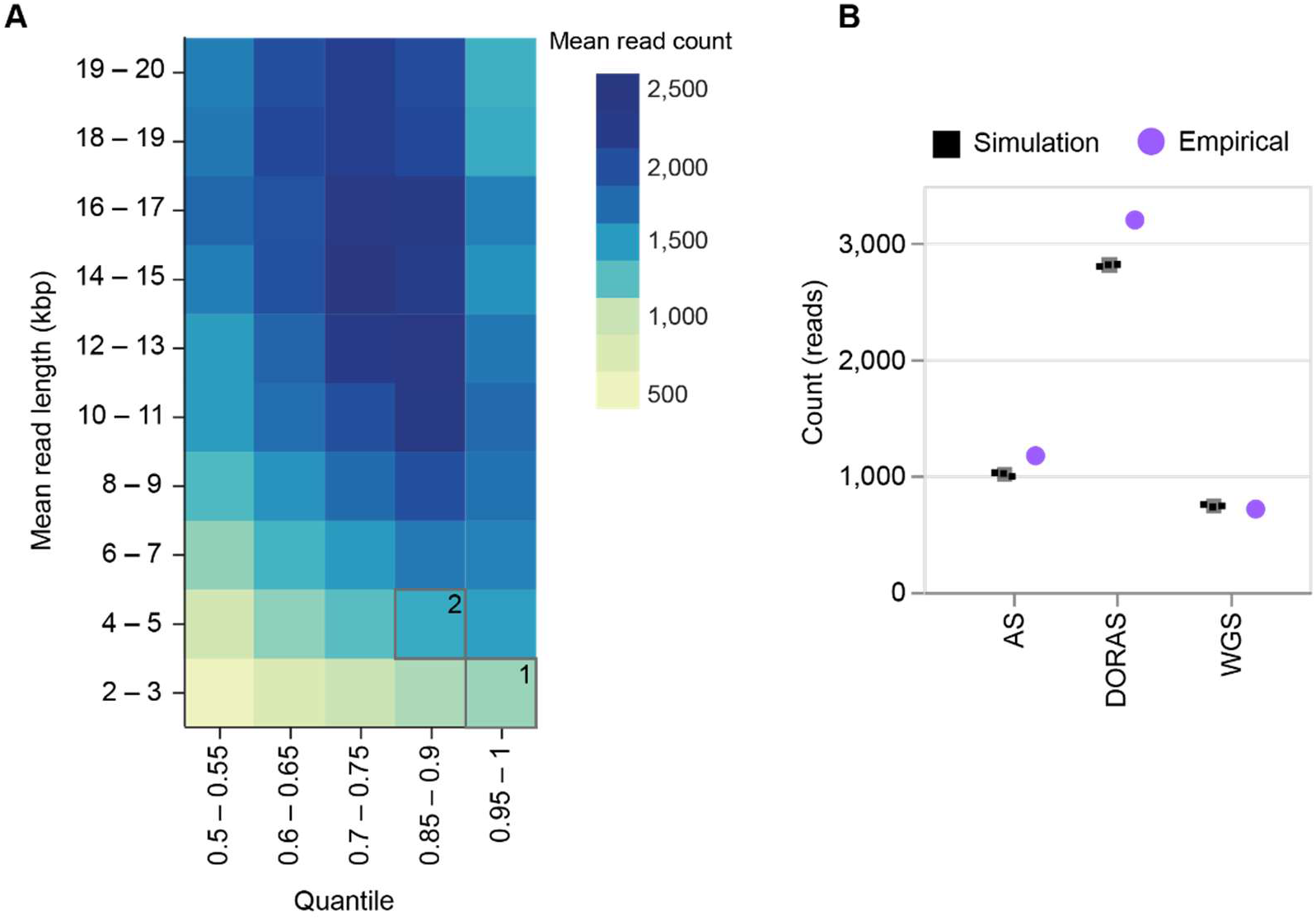
Simulation-driven optimization of AS parameters compared with empirical sequencing runs. **A**) Heat map of AS simulation. The color bar corresponds to the mean count of reads mapping to the ROI for one sample. The horizontal axis depicts the quantile of read length distribution and the vertical axis displays the corresponding read length in kilo base pairs. In black, are highlighted the two experimental read lengths 4.9 kb (^1^non-sheared) and 2.4 kb (^2^sheared). **B**) We simulated three runs for one hour (black squares), one under standard conditions (WGS), one using AS with an arbitrarily extended reference (50 kb reference) and sheared DNA (mean length 2.4 kb) and an optimized version with an optimal reference extension length (18 kb, quantile 85) and non-sheared DNA (DORAS). The simulated runs were compared to real sequencing runs under comparable conditions (purple points).

### AS overperforms WGS with adequate reference context

Building on our results, we designed experiments that compared two conditions: One using sheared DNA fragments with a non-optimized (fixed 25 kb) reference extension, and another using non-sheared (longer) DNA fragments with a reference length optimized according to the model’s optimal length based on the read length distribution (**Fig. 4A**). This allows us to experimentally test how fragment length and reference extension strategy affect AS efficiency and ultimately ROI coverage.

**Figure 4.**
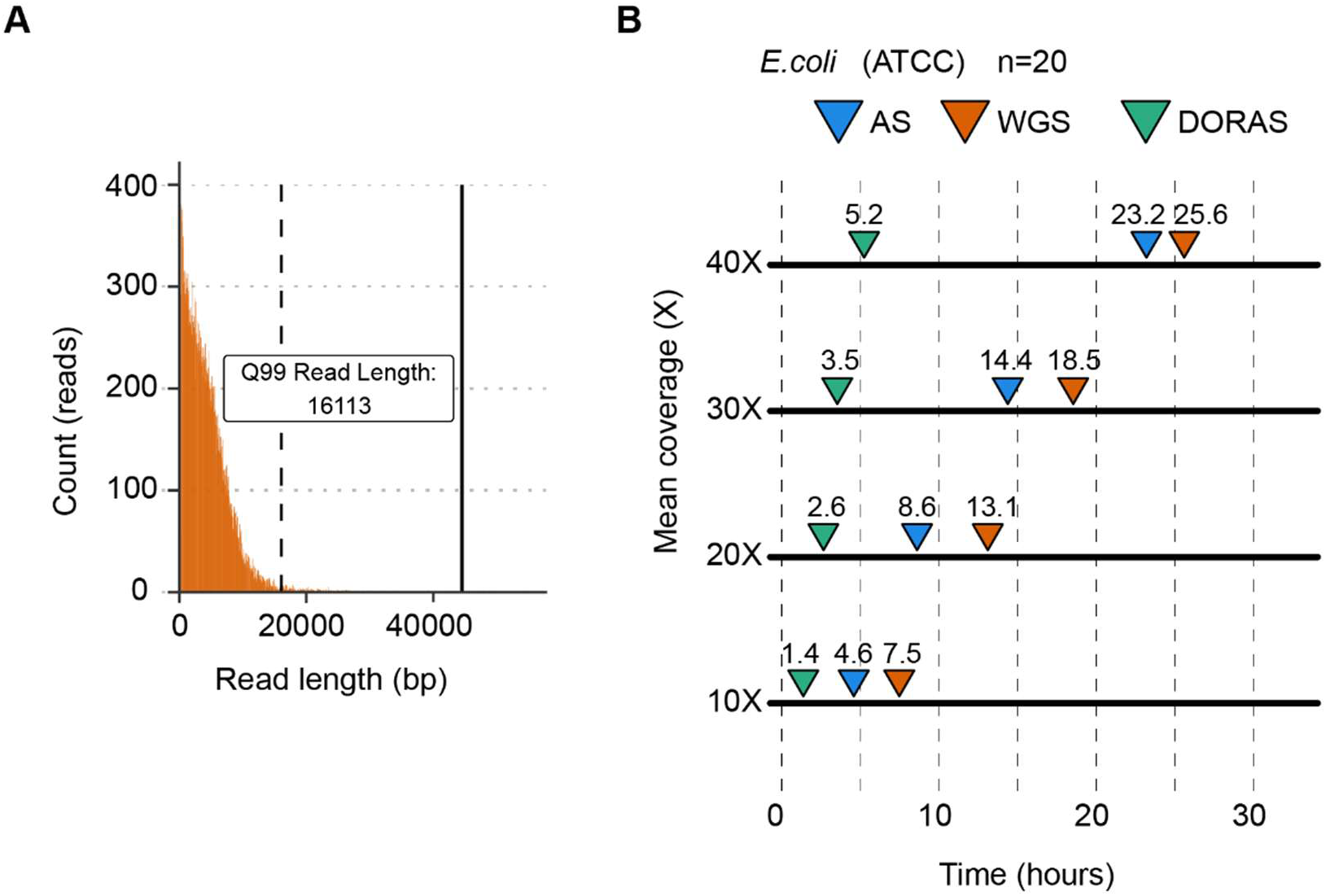
Experimental performance validation of the reference extension with the DORAS protocol. **A)** The optimal reference extension length is defined here as the 99^th^ quantile (dashed line) of read length distribution for the sheared samples. The continuous line defines the maximum value or 100^th^ quantile. **B)** A multiplexed pool of 20 replicates of *E. coli* (ATCC) was sequenced for 72 hours, under standard conditions, or with AS using 50-kb extended reference sequences (25 kb on each ROI side). The downward triangles represent the time when all 20 barcodes reached a specific mean coverage. In blue, gDNA prepared using Covaris shearing, and in green gDNA prepared without shearing. In the DORAS protocol reference extension was optimized according to read size.

We assessed the performance of the approach using 20 barcoded, identically replicated ATCC *E. coli* DNA samples, splitting the library for AS and standard WGS runs. Using a 25-kb extended reference, AS achieved 10X, 20X, 30X, and 40X coverage in 4.6, 8.6, 14.4, and 23.2 hours, respectively, compared to 7.5, 13.1, 18.5, and 25.6 h for standard WGS sequencing (**Fig. 4B, Table S2**). Then, to assess the impact of reference extension and DNA fragment size, we compared the DORAS optimized strategy (q85-based extension with long DNA see **Fig. 3A**) to the non-optimized approach (fixed 25 kb extension with sheared DNA). When both read length (mean length of 4.4 kb) and reference size (18-kb extension) were optimized on the same 20 multiplexed ATCC samples, coverage times were further reduced: 10X, 20X, 30X, and 40X were reached in just 1.4, 2.6, 3.5, and 5.2 h, respectively, confirming the benefit of a model-based reference extension.

We further validated the optimized approach with clinical isolates of *C. diphtheriae* and vancomycin-resistant *Enterococcus*, showing that AS with optimized extended references (*i*.*e*. DORAS) enables efficient and accurate MLST typing (**Fig. 5; Table S3)**. Using recent evaluations of the variant caller Medaka (27) we opted to use a minimum of average coverage of 20X to build consensus sequences (28), followed by a profiling of the ST through the BIGSdb-Pasteur web API (see Methods; **Table 2**). For *C. diphtheriae* (**Fig. 5A**) using the DORAS protocol, the maximum time to reach 20X was 13 h, vs. 42 h with WGS (no AS), corresponding to average times of 4.9 and 8.4 h for DORAS and WGS, respectively. For VRE samples (**Fig. 5B**), the 20X mark was reached after 8 h, whereas 12.8 h under WGS conditions, with mean times to reach 20X of 3.5 and 5.6 h for DORAS and WGS, respectively. Overall, applying DORAS that optimizes the reference size combined with using long DNA molecules markedly accelerates MLST typing for clinical isolates. Across all our evaluations with the three bacterial species considered in our study, 20X coverage was sufficient to obtain 100% accurate consensus sequences (data not shown), as suggested previously (28).

**Table 2.** Sequencing time required to reach a specific coverage.

| Species | Mean coverage | Maximum time (h)<sup>1</sup> |  | Mean time (h)<sup>3</sup> |  |
| --- | --- | --- | --- | --- | --- |
|  |  | WGS | DORAS | WGS | DORAS |
| *C. diphtheriae* | 10X | 24.5 | 7.5 | 8.4 | 2.8 |
|  | 20X | 42.3 | 13.0 | 17.7 | 4.9 |
|  | 30X | 60.8 | 21.2 | 26.5 | 6.8 |
|  | 40X | >72 | 29.0 | >72 | 8.9 |
| *E. faecium* | 10X | 8.8 [n=19] | 5.8 [n=18]<sup>2</sup> | 2.9 | 2.1 |
|  | 20X | 12.8 [n=19] | 8.0 [n=18] | 5.6 | 3.5 |
|  | 30X | 22.1 [n=19] | 11.2 [n=18] | 8.6 | 4.7 |
|  | 40X | 32.8 [n=19] | 14.0 [n=18] | 12.3 | 5.5 |

<sup>1</sup>The maximum time corresponds to the time needed until all samples (batch of 20 isolates) reach the indicated mean coverage. <sup>2</sup>Some VRE isolates did not reach a minimum coverage of 40X after 72 h, and were excluded from the time evaluation. <sup>3</sup>The mean time is the average time across all samples needed to reach the specific mean coverage threshold.

**Figure 5.**
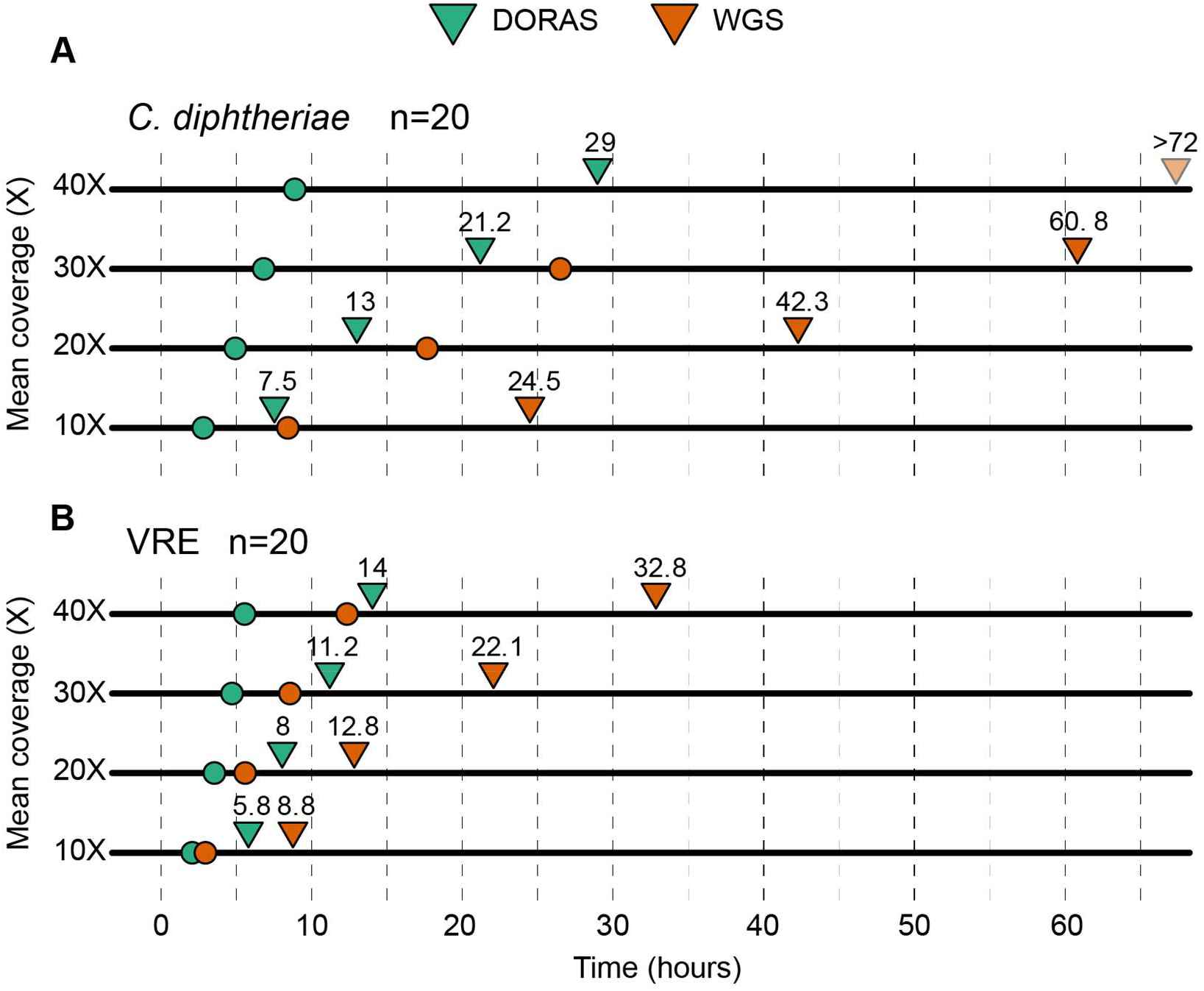
DORAS validation on clinical outbreak isolates. Multiplexed ONT libraries consisting of 20 clinical isolates of **A**) *C. diphtheriae*, and **B**) vancomycin-resistant *E. faecium* (VRE). Downward triangles represent the total sequencing time required for all 20 barcodes to achieve a specific mean coverage, whereas colored circles denote the average time to reach that threshold. Performance across all conditions was benchmarked against standard WGS for the same libraries (orange).

### Leveraging live-generated ONT data to optimally extend the ROI context

Finally, to assess whether our complete workflow enhances MLST gene sequencing under routine conditions, we applied the full DORAS workflow to sequence a panel of 19 clinical *E. coli* isolates obtained during standard diagnostic procedures, along with our ATCC *E. coli* reference strain. We compared the time required to reach specific coverage milestones (10X to 40X) under DORAS conditions (optimized live-generated reference sequences) and standard WGS **(Fig. 6A, Table S4)**. DORAS demonstrated important time gains compared to standard WGS with maximum times to reach specific coverage milestones of 7.44 h (10X), 9.57 h (20X), 13.4 h (30X), and 15.9 h (40X) under DORAS, versus 30.0 h (10X), 30.6 h (20X), and >30 h (for 30 and 40X; the coverage was never reached within the sequencing run duration) under standard WGS. The corresponding mean times to reach these coverages were 2.89, 4.93, 6.49, and 7.66 h for DORAS, compared to 4.95 and 8.67 h for WGS, respectively. These results demonstrate that DORAS achieves the same or higher MLST ROI coverage approximately three to four times faster than conventional WGS for clinical isolates, confirming the practical diagnostic utility of the optimized reference extension strategy.

**Figure 6.**
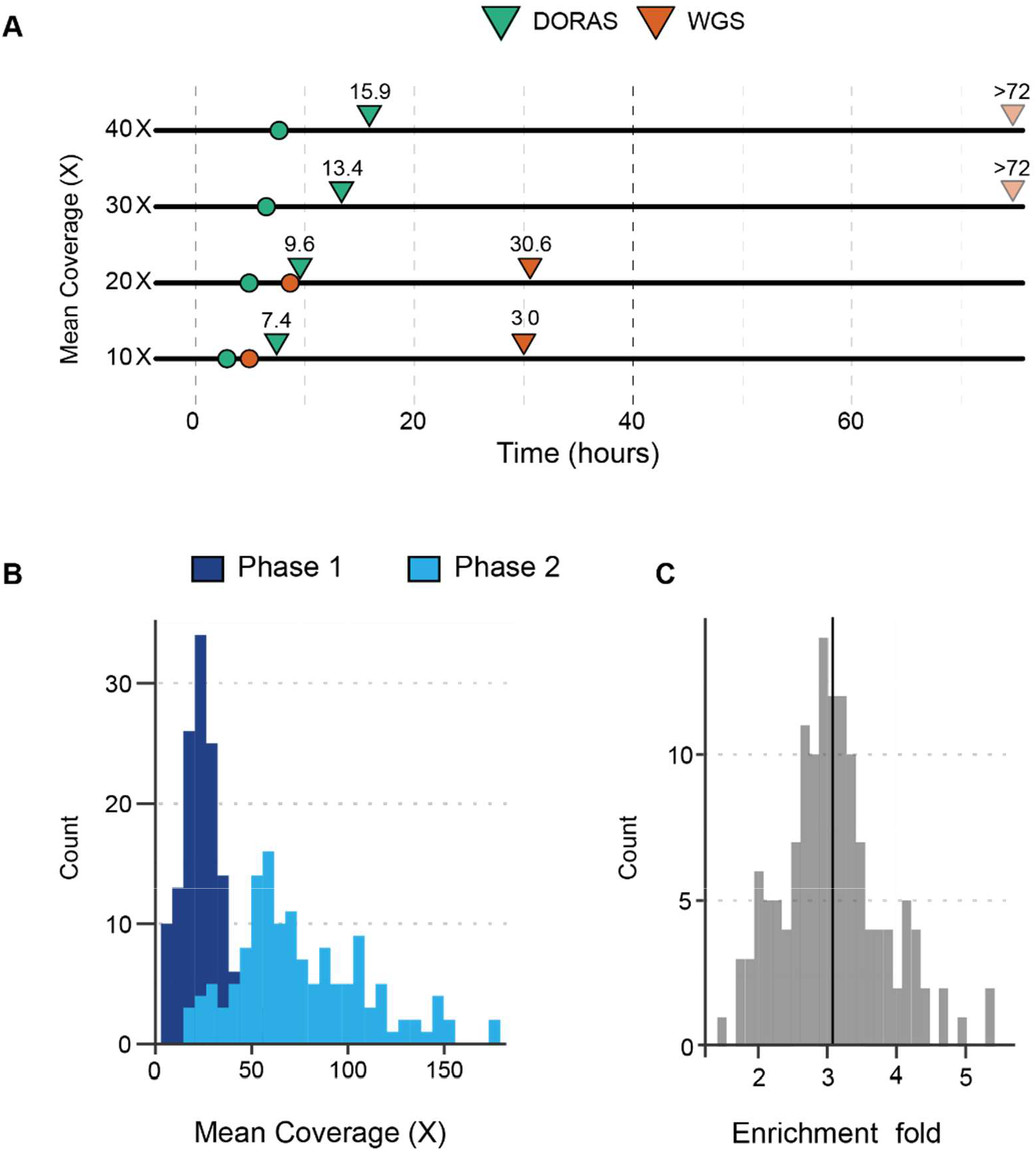
Real-time validation of the DORAS pipeline using clinical *E. coli* samples. **A**) Multiplexed ONT libraries consisting of 19 clinical isolates and one ATCC *E. coli*. Downward triangles and colored circles represent the time when all 20 barcodes reached a specific mean coverage (10X, 20X, 30X and 40X), and colored circles represent the average time to reach this threshold. All sequencing results were ultimately compared to the same library sequenced under standard WGS conditions on a separate flow cell in parallel (orange). **B**) The mean coverage after six sequencing hours was compared between Phase 1 (extension phase, no enrichment) and Phase 2 of DORAS (enrichment, using the live extended reference). **C**) The fold enrichment of the standard sequencing was calculated as the ratio of the maximum mean coverage between phase 1 and phase 2 on the same flow cell after six hours of sequencing.

Additionally, the average mean coverage of our method was evaluated. To do so, we compared the mean coverage reached after six hours of phase 1 (extension) to that reached after six hours of the second phase (query). Mean coverage was strongly increased in phase 2, confirming successful AS read recruitment using the live-generated reference. The mean coverage under standard conditions (WGS) displayed a maximum mean coverage of 25.7X (sd 12.4), and under AS conditions in phase 2, the mean coverage was 75.7X (sd 33.8) (**Fig. 6B**), corresponding to an average fold enrichment of 3.1 (sd 0.7) (**Fig. 6C**). Our results demonstrate that AS with dynamically extended references (DORAS) provides sufficient enrichment of target sequences, and maintains high-quality data acquisition while significantly improving read enrichment efficiency.

### Under high-throughput conditions, WGS provides insufficient coverage for MLST genes

ONT sequencing is well suited to cost-efficient microbial genomics because sample batching can be adjusted flexibly and sequencing runs can be stopped once sufficient data have been generated. In a diagnostic microbiology setting, completing sequencing within approximately 10 hours is particularly advantageous, as it enables an overnight run within a two-day workflow from culture plates to report. We therefore compared WGS and DORAS in a high-throughput *E. coli* multiplexing simulation designed to evaluate the maximal sample load compatible with reliable MLST gene recovery. Specifically, we determined how many samples WGS could process within a fixed 10-hour, 4.2-Gb sequencing window before the least-covered ROI fell below the 20X threshold required for confident sequence-type identification and variant calling. This threshold corresponds to the coverage reached by DORAS within the same time frame (**Fig. 6A**). The DORAS workflow was modeled as a 6-hour extension phase (~3 Gb), followed by a 1.2-Gb query phase in which reads were accepted or rejected according to real-time classification probabilities derived from the extended-reference model. To account for uneven barcode and ROI representation in high-plex nanopore sequencing, multiplexing effects across N barcodes and ROIs were simulated using multinomial sampling with Dirichlet priors and a concentration parameter of 5.

Because read allocation across multiplexed samples and ROIs is stochastic, results are reported as the mean of five simulations (see Methods; **Fig. 7**). As expected, mean sequencing depth per ROI decreased progressively as the number of samples increased, reaching 51.2 reads (sd 26.2) for WGS and 119.5 reads (sd 49.1) for DORAS at 20 samples. The limiting metric was the lowest-depth ROI across all samples, represented by triangles in **Fig. 7**. For WGS, this minimum-depth ROI fell below the 20X threshold when more than 10 samples were multiplexed. In contrast, DORAS maintained the minimum ROI depth above 20X across the full range of tested batch sizes. At 20 barcodes, the lowest-depth ROI yielded approximately 5.2-fold more reads with DORAS than with WGS under equivalent sequencing conditions. Across simulations, the mean enrichment for the lowest-depth ROI was 3.9-fold (sd 1.5). These results identify the multiplexing threshold at which WGS no longer provides sufficient MLST gene coverage and demonstrate that DORAS preserves reliable target recovery under higher-throughput conditions.

**Figure 7.**
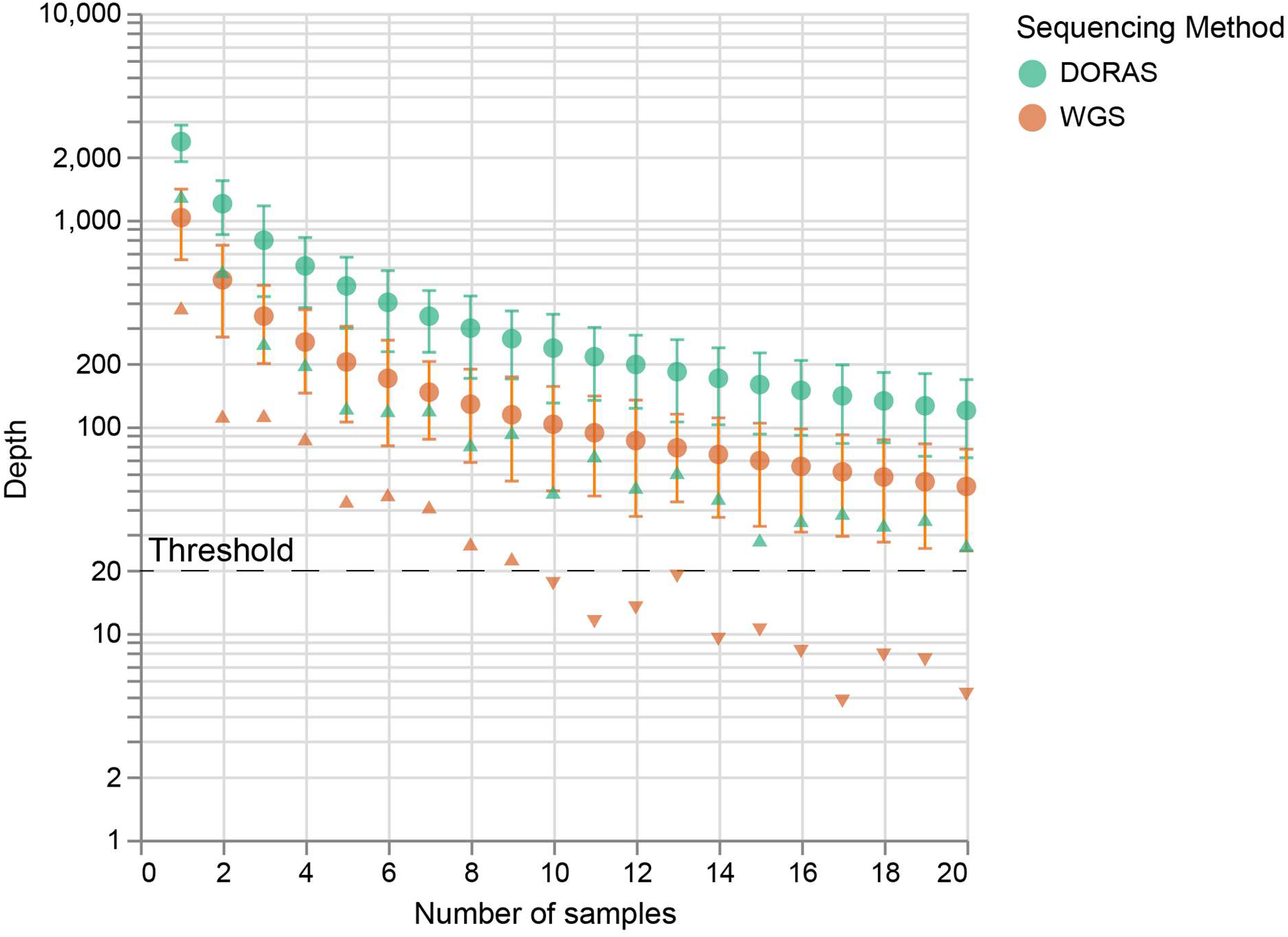
Multiplexing effects in WGS and DORAS. Results of five simulation of varying batch sizes (1 to 20 samples) under WGS (orange) and DORAS (green) conditions, with seven ROIs. The total of giga bases (GB) used was kept constant and the number of reads mapping to the ROI were assigned according to the AS model (see Method section). The circles represent the mean number of reads mapping to the ROI for each batch size and settings. Error bars correspond to one standard deviation. Each triangle represents the minimum coverage obtained among 20 samples and seven ROIs. The triangles facing upwards and downwards correspond to values above and below the 20X threshold, respectively, which was used to call the consensus sequences.

## DISCUSSION

We developed DORAS, a novel workflow designed to overcome the critical limitations of standard AS when applied to small, targeted genomic regions like those used in MLST profiling. For bacteria genomics, this is especially important because genome structure and gene synteny may vary across species and strains (29). Our initial assessment confirmed that conventional ONT AS, using only the short MLST loci as reference sequences, results in a high rejection rate of relevant reads (approximately 79% rejection of target-containing reads). This inefficiency arises when the target regions often fall outside the initial detection window of the sequencing software, leading to ineffective enrichment. Consequently, standard AS performed poorly compared to WGS in achieving sufficient target coverage. By introducing the two-phase DORAS approach, comprising an extension phase to dynamically determine the genomic context of the ROI and a query phase to leverage AS, we successfully enabled efficient enrichment without requiring prior knowledge of the ROI flanking sequences. This streamlined process dramatically reduced the time-to-result compared to standard WGS. Specifically, DORAS achieved full Sequence Type (ST) determination for clinical isolates of *C. diphtheriae* and vancomycin-resistant *Enterococcus* in approximately 3 to 8 hours. We demonstrated that the newly developed workflow to dynamically extend a reference of unknown bacteria allows us to provide enrichment of MLST *loci* to reach an average coverage of 20X within less than 10 hours using a batch of 20 clinical *E. coli* isolates.

The primary novelty of DORAS lies in its dynamic reference extension strategy. While AS is highly effective for enriching large targets like entire plasmids or large eukaryotic genes, its application to multiple, short, non-contiguous targets like MLST loci presents a unique challenge. Our approach is distinct in that it optimizes the reference sequence itself, rather than relying on external software to finely control the rejection process, as done in other tools like readfish or BOSS-RUNS (7, 10). We developed a mathematical model based on sample’s empirical read length distribution and genome size to calculate an optimal reference extension size. This optimization step is crucial: As shown in our validation experiments, using a sub-optimal fixed extension (e.g., 50 kb) on short, sheared DNA resulted in significantly less efficient enrichment compared to the DORAS protocol applied to non-sheared DNA with its calculated extension. As shown in this study, the naïve use of a fixed quantile (*e*.*g*. q99) would hurt AS performances considering the possible variations in library preparation conditions, number and type of reference sequences, and their respective sizes. This dynamic calculation ensures a run-agnostic calculation of the ideal reference to encompass the majority of target-containing reads while minimizing the sequencing of non-informative flanking regions. A principal advantage of the DORAS workflow is that it eliminates the need for PCR amplification. By removing this step, the protocol significantly reduces hands-on time, lowers library preparation cost, and crucially, avoids the inherent amplification biases that often complicate traditional sequence typing methods. The successful validation using several clinically relevant bacterial pathogens and batch sizes confirms the robustness and versatility of the method in routine settings encountered in diagnostic laboratories. This study identifies a definitive multiplexing threshold beyond which WGS is no longer adequate for reliable MLST resolution, whereas DORAS maintains consistent sensitivity even with large batch sizes. Specifically, we observed that WGS performance falls below the requisite detection threshold once multiplexing exceeds 10 samples. This crossover point establishes a practical limit for WGS in high-throughput settings, providing a clear decision framework for selecting DORAS. Consequently, DORAS emerges as an optimal, cost-effective strategy for routine clinical surveillance of large isolate batches or when rapid, targeted typing is required for outbreak investigations.

Despite these significant advances, the current implementation of DORAS has inherent limitations based on its reliance on the native ONT AS function. When sequencing a multiplexed batch of samples, two primary bottlenecks currently constrain the overall time efficiency. First, the Extension Phase requires producing sufficient coverage for the slowest-performing sample to generate extended references. Second, during the Query Phase, the run termination is governed by the sequencing of the slowest-performing sample, which prevents individual barcodes from being stopped as soon as they reach target coverage. Consequently, sequencing resources continue to be consumed by samples that have already been successfully typed. Furthermore, the time-to-result was validated across three clinically relevant bacterial species, including live dynamic reference extension during full sequencing runs. We also evaluated the potential usage of more than seven target genes (ROI) and we concluded that in this current state no more than 100 genes of 500 bp could benefit from the enrichment provided by DORAS (data not shown). This currently excludes the usage of DORAS for core genome MLST (cgMLST) applications where >1,500 ROI are typically used for typing (13). We also evaluated that the total length of the reference sequences used with our current method remains below the 10% cut-off of the genome size recommended by nanopore (23).

Future work may explore integrating the DORAS dynamic reference logic with external adaptive sampling control tools such as readfish (7). While such integration could, in principle, enable dynamic monitoring and termination of the Extension Phase upon reaching coverage thresholds, per-barcode termination during the Query Phase, and a fully automated transition between these phases, it is not straightforward. In practice, this would require tightly coordinating our existing scripts and control logic with external tooling and with the Nanopore built-in adaptive sampling software, which imposes additional architectural and implementation constraints. Nonetheless, if these challenges can be addressed, such an integration could further reduce the overall time-to-result and move towards a fully adaptive, closed-loop system for clinical microbial typing. The DORAS concept, which dynamically defines reference context based on real-time molecular data, could potentially be broadly applicable to other targeted gene sets beyond MLST, such as antimicrobial resistance genes or virulence factors, promising a paradigm shift toward rapid, flexible, and efficient targeted sequencing.

## ACKNOWLEDGEMENTS

The authors thank the diagnostic department of the Institute for Infectious Diseases, University of Bern, for collecting and storing the clinical isolates, and Stefan Neuenschwander with beta testing the DORAS script.

