## Supplementary text, Figures and Tables for "PCR-free, targeted genomic sequencing using Dynamically optimized reference Adaptive Sampling (DORAS)"

**1 Supplementary Text** ... 2

1.1 Adaptive sampling model ... 2

1.1.1 Model the read length distribution ... 2

1.1.2 Simulating a library of read lengths ... 3

1.1.3 Size of the extended reference ... 3

1.1.4 Total number of k-mers in a *single* haploid genome ... 3

1.1.5 How many k-mers map **only** to the *ROI itself* ... 4

1.1.6 Distribution of **k-mers** in the *extended* reference (ROI + flanking sequence) 4

1.1.7 Special handling of **very long reads** ... 4

1.1.8 Ratios that describe the *distribution of reads* ... 4

1.2 Time simulation of AS ... 5

*1.2.1 Notation* ... 5

1.2.2 Pore decay ... 6

1.2.3 Event at a free pore ... 6

1.2.4 Update of a busy pore (i.e. currently sequencing) ... 6

1.2.5 Whole-run recursion ... 6

**2 Supplementary Figures** ... 8

**3 Supplementary Tables** ... 10

### 1 Supplementary Text

In this section, we provide additional information for reader of the DORAS manuscript. First, we describe the mathematical model used to understand the intricate relationships between the length of the reference and the length of the reads. We proceed with additional details about the main figures and also provide tables of the corresponding samples used in each figure presented in the manuscript.

#### 1.1 Adaptive sampling model

The adaptive-sampling (AS) process relies on the simple principle of enrichment (or exclusion) of reads that map to a provided reference within the first 500 bp when entering the pore. The number of reads detected is binomial $y \sim Bin(\theta; n)$ (n corresponds to number of reads generated). Here $\theta$ represents the probability of a read (here modeled by k-mers) to contain a mappable portion (to the provided ROI) within the first 500 bp of the read. In this study, we focus on the enrichment of small ROI within a bacterial genome that are small enough to be completely engulfed in a read outside of detectable region of 500 bp at the beginning of the read. In order to increase the chance of such a region to be detected it is necessary to increase the flanking regions of this ROI (buffer region in the nanopore documentation). In the context of nanopore sequencing reads are not of constant length. Thus, we also aim to take into account the variability in read length and propagate this to the computation of the ideal extension. Finally, the probability $\theta$ of a detecting (to enrich) can be estimated as $(\hat{\theta})$ as the ratio of all possible k-mers contained in the extended reference over the total k-mers contained in a genome.

Below we outline the three logical steps that turn a purely combinatorial count of k-mers into a probabilistic estimate of $\theta$ to answer “will a random read map to the reference?"

##### 1.1.1 Read length distribution modelling

We will first describe how we chose to model the read length and use to obtain the distribution of all combination of k-mers contained in the entire genome and next the ones containing the ROI. From the distribution of read length one can obtain the quantile (e.g. quantile 0.99) of this distribution.

Table of the notation used:

| Symbol (math) | Description |
|---|---|
| $\mu$ | Target *mean* read length (in base pairs) |
| $\sigma$ | Target standard deviation (in base pairs) |
| $k$ | Number of reads to use for the simulation (default $k = 1000$) |
| $q$ | Quantile used to define the “long-read” cut-off (default $0.99$) |
| $c_{\text{ROI}}$ | Number of ROI chosen (default to 7). |

| Symbol (math) | Description |
|---|---|
| $L_{\text{ROI}}$ | Length (in bp) of the ROI, the part we ultimately want to detect (default $500$ bp). |
| $L_{reads}$ | Lengths (in bp) of the individual reads. |
| $m$ | Number of bases removed from each read to account for the two flanking ("buffer") regions that cannot be used for k-mer counting (default $100$ bp). |
| $R_{ext}$ | Extended reference |
| $S_{opt}$ | Size of the *extended* reference $R_{ext}$ that contains the ROI. |

##### 1.1.2 Simulating a library of read lengths

We draw $k$ reads from a *Gamma* distribution whose parameters are chosen so that the *expected* value equals the supplied mean $\mu$ and the *standard deviation* of that mean or from a set of vectors of read length extracted from a FASTQ file.

1. **Shape (α) and scale (θ) parameters** for a Gamma distribution with mean $\mu$ and standard deviation $\sigma$ are

$$\alpha = \frac{\mu^2}{\sigma^2}, \qquad \theta = \frac{\sigma^2}{\mu}.$$

2. **Sample**

$$L_{reads} = L_1, L_2, \ldots, L_k \sim Gamma(\alpha, \theta).$$

##### 1.1.3 Size of the extended reference

The extension of the reference is computed from the given read length distribution $L_{reads}$ by computing the quantile $q$ from the

$$S_{opt} = 2\, Quantile_q\, (L_{reads}) - L_{ROI}.$$

*(Note: In the code implementation, this value can also be arbitrarily set for testing purposes)*

##### 1.1.4 Total number of k-mers in a *single* haploid genome

The total number of distinct k-mers that can be taken from a *complete* genome of length $G$ (where $G$ is the genome size) is

$$N_{\text{total } k\text{-mers}} = G - L_{\text{reads}} + 1$$

##### 1.1.5 How many k-mers map **only** to the *ROI itself*

If a read is *short enough* that it does **not** reach the flanking part of the extended reference (i.e. it lies completely inside the detection window), the number of k-mers that intersect **only** the ROI is

$$K_{\text{ROI-alone}}(L) = c_{\text{ROI}}\ (L_{\text{ROI}} + L - 2\,m).$$

Again this is applied element-wise to the whole vector of read lengths:

$$K_{\text{ROI-alone}} = c_{\text{ROI}}\ (L_{\text{ROI}} + \mathrm{L}_{\text{reads}} - 2\,m).$$

##### 1.1.6 Distribution of k-mers in the *extended* reference (ROI + flanking sequence)

Once the $S_{\text{opt}}$ is known, we estimate the distribution of k-mers belonging to the extended reference. Here we only take into account k-mers that contain the full ROI, and k-mers mapping up to the *m* bp (default 100) of the end of the extended reference $R_{ext}$.

$$K_{\text{ref}}(L) = \mathrm{c}_{\text{ROI}}\left(\mathrm{S}_{\text{opt}} + 2\mathrm{L}_{reads} - 2\mathrm{m} - \mathrm{L}_{reads} + 1\right) = \mathrm{c}_{\text{ROI}}\left(\mathrm{S}_{\text{opt}} + \mathrm{L}_{reads} - 2\mathrm{m} + 1\right)$$

*(Note: When the function works on a whole vector of read lengths this becomes the element-wise array.)*

$$\mathrm{K}_{\text{ref}} = c_{\text{ROI}}\ \left(S_{\text{opt}} + \mathrm{L}_{\text{reads}} - 2\,m + 1\right).$$

##### 1.1.7 Handling of very long reads

A read that is **longer** than the reference *minus* a detection window $\varepsilon$ (set to 500 bp) will *span* the whole reference and there *cannot* be uniquely assigned to the ROI alone.

$$\mathrm{M}_{\text{long}} = \left\{L > S_{\text{opt}} - 2\,\varepsilon\right\}.$$

For those reads the number of ROI-k-mers is **re-computed** with the *detection window* in place of the full reference:

$$K_{\text{ref}}^{(\text{long})}(L) = c_{\text{ROI}}\ \left(2\left(\varepsilon + S_{\text{opt}} - 2\,m\right)\right).$$

##### 1.1.8 Ratios that describe the *distribution of reads*

After establishing the distributions of the combination of k-mers inside $R_{ext}$ and the entire genome $N_{total\ k\text{-}mers}$, we can compute, using the k-mers, an estimation of the probability of a read entering the pore to be enriched. We can compute then:

- $N_{total\ k\text{-}mers} = G - \mathrm{L}_{\text{reads}}\ +\ 1$ (total number of k-mers spanning the genome).

- $\Pi_{ref} = \frac{K_{ref}}{N_{total}}$ , estimated probability of finding a k-mer that maps to the extended ref. This corresponds to the probability of a read being accepted during AS ($\hat{\theta}$ mentioned earlier).

- $\Pi_{\text{ROI alone}} = \frac{\mathbf{K}_{\text{ROI-alone}}}{N_{\text{total}}}$, this is estimated probability of a read mapping to the ROI. This gives the probability of finding a read mapping to the ROI using standard sequencing (WGS).

- $\Pi_{\text{ROI-Ref}} = \frac{\mathbf{K}_{\text{ROI-alone}}}{K_{ref}}$**,** this is the probability of a k-mer mapping to the $K_{ref}$ also maps to the ROI $P(ROI|K_{ref})$. This probability is used at the end of the simulation to estimate how many reads mapping to the $Ref_{ext}$ ($\mathcal{A}$ , accepted molecules see next section 2.1) also map to the ROI.

#### 1.2 Time simulation of AS

Using AS, the decision to accept or reject a read takes place at the pore level. If a read is accepted (mapping to the reference provided), this pore will sequence the entire molecule. To mimic this process, each time a read is accepted the pore is occupied for the time lapse proportional to the length of the read.

##### *1.2.1 Notation*

| Symbol | Meaning |
|---|---|
| $T$ | total simulation time (seconds) |
| $\Delta t = 1$ s | length of one discrete time step |
| $P_{\max}$ | maximum number of pores ( default 500) |
| $P_0$ | initial number of pores at $t = 0$ |
| $L$ | observed fragment length (bases) drawn from $\mathcal{L}$ |
| $\mathcal{L}$ | discrete distribution from which fragment lengths are drawn |
| $\pi_{ref,t}$ | If 1 *new* fragment belongs to the extended reference at second $t$, else 0 does not belong to the extended reference |
| $\Pi_{ref}$ | Probability of read to be accepted (enriched by AS) |
| $\Pi_{\text{alone}}$ | list of ROI probabilities used in WGS mode |
| $\mathbf{s}_t = (s_{t,1}, \dots, s_{t,P_{\max}})$ | vector of *remaining* times for each pore at second $t$; $s_{t,i} \leq 0$ means pore *i* is free, $s_{t,i} > 0$ means it is busy |
| $c_{\text{rej}} = 2$ s | Fixed “rejection” time when a fragment is discarded (non-WGS mode) |
| $v = 400$ bases · s$^{-1}$ | Sequencing speed (the code uses `length/400`) |
| $N_{\text{reads}}$ | Counter of total fragment attempts |
| $\mathcal{A}$ | Count of the number of reads belonging to the extended reference |

##### 1.2.2 Pore decay

The simulation model assumes no decay over time but the number of pores can be adjusted ($P_0$) to simulate the pore behavior of a flow cell.

##### 1.2.3 Event at a free pore

For each pore $i$ such that $s_{t,i} \leq 0$, the algorithm performs:

1. Increment the global read counter

   $N_{\text{reads}} \leftarrow N_{\text{reads}} + 1.$

2. Sample a ROI-probability $\pi_t$

$$\pi_t = \begin{cases} Bin(\Pi, 1) & \text{if enrichment mode ( WGS = False),} \\ Bin(\Pi_{\text{alone}}, 1) & \text{if WGS mode (WGS = True).} \end{cases}$$

3. Verify if accepted or not

   *If* $\pi_t = 1$ (fragment belongs to ref) **accept** the fragment:

$$L \sim \mathcal{L}, \qquad s_{t+1,i} = \frac{L}{v}, \qquad \mathcal{A} \leftarrow \mathcal{A} \cup L.$$

   *Else* (fragment rejected)

$$\begin{cases} \text{if WGS} & L \sim \mathcal{L},\ s_{t+1,i} = L/v \quad \text{(sequencing anyway)} \\ \text{otherwise} & s_{t+1,i} = c_{\text{rej}}. \end{cases}$$

##### 1.2.4 Update of a busy pore (i.e. currently sequencing)

If a pore is busy ($s_{t,i} > 0$) we simply decrement ($\Delta t = 1s$) the remaining time:

$s_{t+1,i} = s_{t,i} - \Delta t.$

When the counter reaches $0$ (or becomes negative) the pore becomes free again at the **next** second. Here we assume that the pore is available directly after the sequencing is finished.

##### 1.2.5 Whole-run recursion

Putting everything together, for each second $t = 0, \dots, T-1$ we have

$$\begin{array}{cc} s_{t,i} & \text{the inital status of each pore} \\ \text{iterate over pores} & \text{for each } i \in 1, \dots, P_{\max}\text{:} \\ \text{if } i > a_t & \text{(busy pore)} \ \Rightarrow \ s_{t+1,i} = s_{t,i} = 0 \text{ (skip)} \\ \text{else if } s_{t,i} \leq 0 & \text{(free pore)} \ \Rightarrow \ \text{perform steps 1–4 above} \\ \text{else} & s_{t+1,i} = s_{t,i} - 1. \end{array}$$

The simulation stops when $t = T$.

The returned quantities are

| | |
|---|---|
| total_reads | $= N_{\text{reads}}$ |
| accepted_molecules | $= \mathcal{A}$ |
| alive_pores | $= a_T$ |

### 2 Supplementary Figures

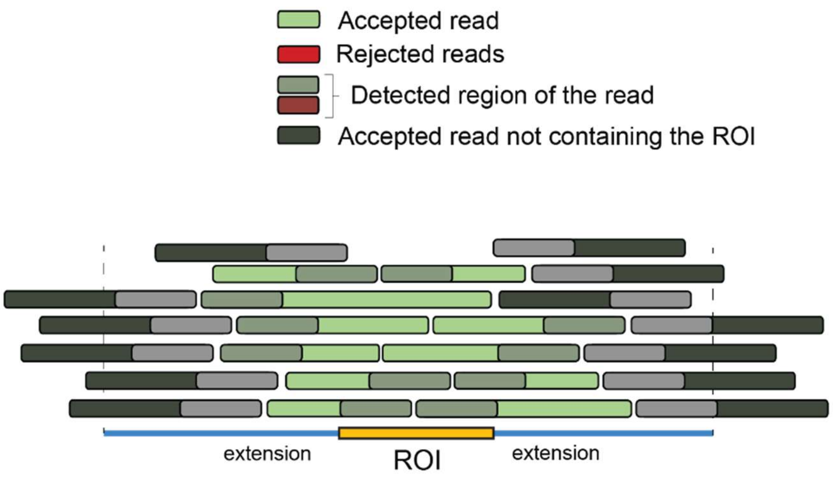


**Figure S1**. Conceptual representation of ROI sequence extension. Overextension is detrimental for efficient AS and ROI read recruitment. Representation of the problem with a reference that is too short where the red reads will not be enriched during AS process.

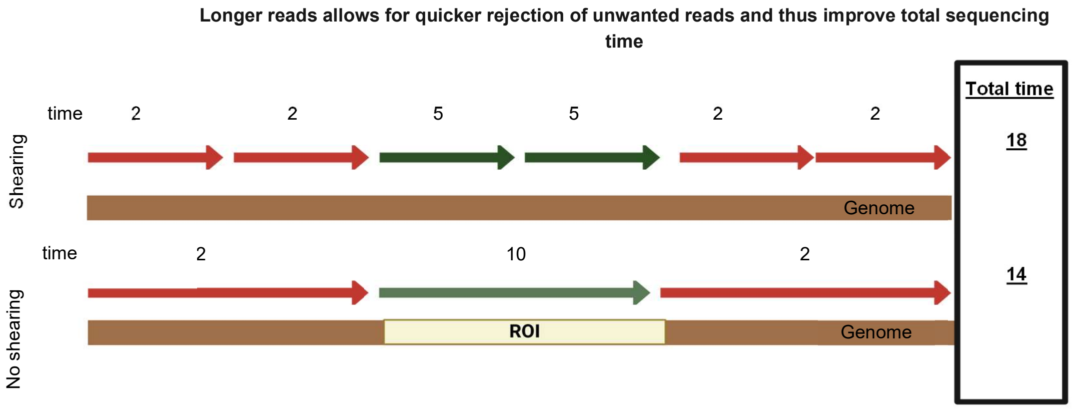


**Figure S2**. Longer reads are advantageous in adaptive sampling. The time needed to sequence fully (green, accepted reads) depends on the size. Oppositely, partial sequencing (red, rejected reads) is independent of the size of the reads which results in a quicker rejection of the unwanted reads and thus quicker sequencing times.

### 3 Supplementary Tables

**Table S1. Obtained Sequence Types (ST) for the ATCC *E. coli* under each sequencing setting, and corresponding accession numbers.**

| Sample | ST | Origin | Settings | NCBI Accession number |
|---|---|---|---|---|
| **sample01** | 1115 | *E. coli* | Standard AS | SAMN56595307 |
| **sample02** | 1115 | *E. coli* | Standard AS | SAMN56595308 |
| **sample03** | 1115 | *E. coli* | Standard AS | SAMN56595309 |
| **sample04** | 1115 | *E. coli* | Standard AS | SAMN56595310 |
| **sample05** | 1115 | *E. coli* | WGS | SAMN56595303 |
| **sample06** | 1115 | *E. coli* | WGS | SAMN56595304 |
| **sample07** | 1115 | *E. coli* | WGS | SAMN56595305 |
| **sample08** | 1115 | *E. coli* | WGS | SAMN56595306 |

**Table S2. Sequence Type of the ATCC *E. coli* for each setting, and their corresponding accession numbers. ST was determined when all samples reached a minimum of 20X.**

| Sample | ST | Origin | Settings | Full resolution time (hours) | NCBI Accession number |
|---|---|---|---|---|---|
| **sample01** | 1115 | *E. coli* | doras | 2.6 | SAMN56595331 |
| **sample02** | 1115 | *E. coli* | doras | 2.6 | SAMN56595332 |
| **sample03** | 1115 | *E. coli* | doras | 2.6 | SAMN56595333 |
| **sample04** | 1115 | *E. coli* | doras | 2.6 | SAMN56595334 |
| **sample05** | 1115 | *E. coli* | doras | 2.6 | SAMN56595335 |
| **sample06** | 1115 | *E. coli* | doras | 2.6 | SAMN56595336 |
| **sample07** | 1115 | *E. coli* | doras | 2.6 | SAMN56595337 |
| **sample08** | 1115 | *E. coli* | doras | 2.6 | SAMN56595338 |
| **sample09** | 1115 | *E. coli* | doras | 2.6 | SAMN56595339 |
| **sample10** | 1115 | *E. coli* | doras | 2.6 | SAMN56595340 |
| **sample11** | 1115 | *E. coli* | doras | 2.6 | SAMN56595341 |
| **sample12** | 1115 | *E. coli* | doras | 2.6 | SAMN56595342 |
| **sample13** | 1115 | *E. coli* | doras | 2.6 | SAMN56595343 |
| **sample14** | 1115 | *E. coli* | doras | 2.6 | SAMN56595344 |
| **sample15** | 1115 | *E. coli* | doras | 2.6 | SAMN56595345 |
| **sample16** | 1115 | *E. coli* | doras | 2.6 | SAMN56595346 |
| **sample17** | 1115 | *E. coli* | doras | 2.6 | SAMN56595347 |
| **sample18** | 1115 | *E. coli* | doras | 2.6 | SAMN56595348 |
| **sample19** | 1115 | *E. coli* | doras | 2.6 | SAMN56595349 |
| **sample20** | 1115 | *E. coli* | doras | 2.6 | SAMN56595350 |
| **sample01** | 1115 | *E. coli* | AS_sheared | 8.6 | SAMN56595351 |
| **sample02** | 1115 | *E. coli* | AS_sheared | 8.6 | SAMN56595352 |
| **sample03** | 1115 | *E. coli* | AS_sheared | 8.6 | SAMN56595353 |
| **sample04** | 1115 | *E. coli* | AS_sheared | 8.6 | SAMN56595354 |
| **sample05** | 1115 | *E. coli* | AS_sheared | 8.6 | SAMN56595355 |
| **sample06** | 1115 | *E. coli* | AS_sheared | 8.6 | SAMN56595356 |
| **sample07** | 1115 | *E. coli* | AS_sheared | 8.6 | SAMN56595357 |
| **sample08** | 1115 | *E. coli* | AS_sheared | 8.6 | SAMN56595358 |
| **sample09** | 1115 | *E. coli* | AS_sheared | 8.6 | SAMN56595359 |
| **sample10** | 1115 | *E. coli* | AS_sheared | 8.6 | SAMN56595360 |
| **sample11** | 1115 | *E. coli* | AS_sheared | 8.6 | SAMN56595361 |

**Table S2. Sequence Type of the ATCC *E. coli* for each setting, and their corresponding accession numbers. ST was determined when all samples reached a minimum of 20X.**

| Sample | ST | Origin | Settings | Full resolution time (hours) | NCBI Accession number |
|---|---|---|---|---|---|
| **sample12** | 1115 | *E. coli* | AS_sheared | 8.6 | SAMN56595362 |
| **sample13** | 1115 | *E. coli* | AS_sheared | 8.6 | SAMN56595363 |
| **sample14** | 1115 | *E. coli* | AS_sheared | 8.6 | SAMN56595364 |
| **sample15** | 1115 | *E. coli* | AS_sheared | 8.6 | SAMN56595365 |
| **sample16** | 1115 | *E. coli* | AS_sheared | 8.6 | SAMN56595366 |
| **sample17** | 1115 | *E. coli* | AS_sheared | 8.6 | SAMN56595367 |
| **sample18** | 1115 | *E. coli* | AS_sheared | 8.6 | SAMN56595368 |
| **sample19** | 1115 | *E. coli* | AS_sheared | 8.6 | SAMN56595369 |
| **sample20** | 1115 | *E. coli* | AS_sheared | 8.6 | SAMN56595370 |
| **sample01** | 1115 | *E. coli* | WGS | 13.1 | SAMN56595523 |
| **sample02** | 1115 | *E. coli* | WGS | 13.1 | SAMN56595524 |
| **sample03** | 1115 | *E. coli* | WGS | 13.1 | SAMN56595525 |
| **sample04** | 1115 | *E. coli* | WGS | 13.1 | SAMN56595526 |
| **sample05** | 1115 | *E. coli* | WGS | 13.1 | SAMN56595527 |
| **sample06** | 1115 | *E. coli* | WGS | 13.1 | SAMN56595528 |
| **sample07** | 1115 | *E. coli* | WGS | 13.1 | SAMN56595529 |
| **sample08** | 1115 | *E. coli* | WGS | 13.1 | SAMN56595530 |
| **sample09** | 1115 | *E. coli* | WGS | 13.1 | SAMN56595531 |
| **sample10** | 1115 | *E. coli* | WGS | 13.1 | SAMN56595532 |
| **sample11** | 1115 | *E. coli* | WGS | 13.1 | SAMN56595533 |
| **sample12** | 1115 | *E. coli* | WGS | 13.1 | SAMN56595534 |
| **sample13** | 1115 | *E. coli* | WGS | 13.1 | SAMN56595535 |
| **sample14** | 1115 | *E. coli* | WGS | 13.1 | SAMN56595536 |
| **sample15** | 1115 | *E. coli* | WGS | 13.1 | SAMN56595537 |
| **sample16** | 1115 | *E. coli* | WGS | 13.1 | SAMN56595538 |
| **sample17** | 1115 | *E. coli* | WGS | 13.1 | SAMN56595539 |
| **sample18** | 1115 | *E. coli* | WGS | 13.1 | SAMN56595540 |
| **sample19** | 1115 | *E. coli* | WGS | 13.1 | SAMN56595541 |
| **sample20** | 1115 | *E. coli* | WGS | 13.1 | SAMN56595542 |

**Table S3. Sequence Types (ST) of all the samples indicated were verified at 8 (*C. diphtheriae*) and 13 hours (VRE). The samples associated with too few reads are indicated with "-" for the ST and resolution time. They did not reach the threshold of 20X within the sequencing run duration. Accession numbers for both phase 1 and phase 2 are displayed in the case of a DORAS experiments, otherwise only those for phase 1 (WGS) are displayed.**

| Sample | ST | Origin | Settings | Full resolution time (hours) | NCBI Accession number (Phase1, Phase2) | |
|---|---|---|---|---|---|---|
| **sample01** | 384 | CDIP | DORAS | 13 | SAMN56595391 | SAMN56595411 |
| **sample02** | 384 | CDIP | DORAS | 13 | SAMN56595392 | SAMN56595412 |
| **sample03** | 377 | CDIP | DORAS | 13 | SAMN56595393 | SAMN56595413 |
| **sample04** | 384 | CDIP | DORAS | 13 | SAMN56595394 | SAMN56595414 |
| **sample05** | 377 | CDIP | DORAS | 13 | SAMN56595395 | SAMN56595415 |
| **sample06** | 384 | CDIP | DORAS | 13 | SAMN56595396 | SAMN56595416 |
| **sample07** | 698 | CDIP | DORAS | 13 | SAMN56595397 | SAMN56595417 |
| **sample08** | 377 | CDIP | DORAS | 13 | SAMN56595398 | SAMN56595418 |
| **sample09** | 384 | CDIP | DORAS | 13 | SAMN56595399 | SAMN56595419 |
| **sample10** | 698 | CDIP | DORAS | 13 | SAMN56595400 | SAMN56595420 |
| **sample11** | 698 | CDIP | DORAS | 13 | SAMN56595401 | SAMN56595421 |
| **sample12** | 377 | CDIP | DORAS | 13 | SAMN56595402 | SAMN56595422 |
| **sample13** | 377 | CDIP | DORAS | 13 | SAMN56595403 | SAMN56595423 |
| **sample14** | 574 | CDIP | DORAS | 13 | SAMN56595404 | SAMN56595424 |
| **sample15** | 574 | CDIP | DORAS | 13 | SAMN56595405 | SAMN56595425 |
| **sample16** | 377 | CDIP | DORAS | 13 | SAMN56595406 | SAMN56595426 |
| **sample17** | 377 | CDIP | DORAS | 13 | SAMN56595407 | SAMN56595427 |
| **sample18** | 103 | CDIP | DORAS | 13 | SAMN56595408 | SAMN56595428 |
| **sample19** | 377 | CDIP | DORAS | 13 | SAMN56595409 | SAMN56595429 |
| **sample20** | 574 | CDIP | DORAS | 13 | SAMN56595410 | SAMN56595430 |
| **sample01** | 796 | VRE | DORAS | 8 | SAMN56595451 | SAMN56595431 |
| **sample02** | 78 | VRE | DORAS | 8 | SAMN56595452 | SAMN56595432 |
| **sample03** | 796 | VRE | DORAS | 8 | SAMN56595453 | SAMN56595433 |
| **sample04** | 117 | VRE | DORAS | 8 | SAMN56595454 | SAMN56595434 |
| **sample05** | 117 | VRE | DORAS | 8 | SAMN56595455 | SAMN56595435 |
| **sample06** | 117 | VRE | DORAS | 8 | SAMN56595456 | SAMN56595436 |
| **sample08** | 117 | VRE | DORAS | 8 | SAMN56595457 | SAMN56595437 |

**Table S3. Sequence Types (ST) of all the samples indicated were verified at 8 (*C. diphtheriae*) and 13 hours (VRE). The samples associated with too few reads are indicated with "-" for the ST and resolution time. They did not reach the threshold of 20X within the sequencing run duration. Accession numbers for both phase 1 and phase 2 are displayed in the case of a DORAS experiments, otherwise only those for phase 1 (WGS) are displayed.**

| Sample | ST | Origin | Settings | Full resolution time (hours) | NCBI Accession number (Phase1, Phase2) | |
|---|---|---|---|---|---|---|
| **sample07** | 796 | VRE | DORAS | 8 | SAMN56595458 | SAMN56595438 |
| **sample08** | - | VRE | DORAS | - | SAMN56595459 | SAMN56595439 |
| **sample09** | 117 | VRE | DORAS | 8 | SAMN56595460 | SAMN56595440 |
| **sample10** | 117 | VRE | DORAS | 8 | SAMN56595461 | SAMN56595441 |
| **sample11** | 117 | VRE | DORAS | 8 | SAMN56595462 | SAMN56595442 |
| **sample12** | 117 | VRE | DORAS | 8 | SAMN56595463 | SAMN56595443 |
| **sample13** | 796 | VRE | DORAS | 8 | SAMN56595464 | SAMN56595444 |
| **sample14** | 117 | VRE | DORAS | 8 | SAMN56595465 | SAMN56595445 |
| **sample15** | 133 | VRE | DORAS | 8 | SAMN56595466 | SAMN56595446 |
| **sample16** | 80 | VRE | DORAS | 8 | SAMN56595467 | SAMN56595447 |
| **sample18** | 117 | VRE | DORAS | 8 | SAMN56595468 | SAMN56595448 |
| **sample19** | 117 | VRE | DORAS | 8 | SAMN56595469 | SAMN56595449 |
| **sample20** | - | VRE | DORAS | - | SAMN56595470 | SAMN56595450 |

**Table S4. Sequence types for routine clinical *E. coli*. The samples containing too few reads were excluded since they did not reach the threshold of 20X. Accession numbers for both phase 1 and phase 2 are displayed in the case of a DORAS experiments, otherwise only those for phase 1 (WGS) are displayed.**

| Sample | ST | Origin | Settings | Full resolution time (hours) | NCBI Accession number (Phase1, Phase2) | |
|---|---|---|---|---|---|---|
| **sample01** | 1115 | *E. coli* | DORAS | 10 | SAMN56595483 | SAMN56595503 |
| **sample02** | 69 | *E. coli* | DORAS | 10 | SAMN56595484 | SAMN56595504 |
| **sample03** | 131 | *E. coli* | DORAS | 10 | SAMN56595485 | SAMN56595505 |
| **sample04** | 12 | *E. coli* | DORAS | 10 | SAMN56595486 | SAMN56595506 |
| **sample05** | 155 | *E. coli* | DORAS | 10 | SAMN56595487 | SAMN56595507 |
| **sample06** | 10 | *E. coli* | DORAS | 10 | SAMN56595488 | SAMN56595508 |
| **sample07** | 95 | *E. coli* | DORAS | 10 | SAMN56595489 | SAMN56595509 |
| **sample08** | 131 | *E. coli* | DORAS | 10 | SAMN56595490 | SAMN56595510 |
| **sample09** | 15061 | *E. coli* | DORAS | 10 | SAMN56595491 | SAMN56595511 |
| **sample10** | 69 | *E. coli* | DORAS | 10 | SAMN56595492 | SAMN56595512 |
| **sample11** | 69 | *E. coli* | DORAS | 10 | SAMN56595493 | SAMN56595513 |
| **sample12** | 73 | *E. coli* | DORAS | 10 | SAMN56595494 | SAMN56595514 |
| **sample13** | 117 | *E. coli* | DORAS | 10 | SAMN56595495 | SAMN56595515 |
| **sample14** | 538 | *E. coli* | DORAS | 10 | SAMN56595496 | SAMN56595516 |
| **sample15** | 5640 | *E. coli* | DORAS | 10 | SAMN56595497 | SAMN56595517 |
| **sample16** | 567 | *E. coli* | DORAS | 10 | SAMN56595498 | SAMN56595518 |
| **sample17** | 69 | *E. coli* | DORAS | 10 | SAMN56595499 | SAMN56595519 |
| **sample18** | 131 | *E. coli* | DORAS | 10 | SAMN56595500 | SAMN56595520 |
| **sample19** | 131 | *E. coli* | DORAS | 10 | SAMN56595501 | SAMN56595521 |
| **sample20** | 1115 | *E. coli* | DORAS | 10 | SAMN56595502 | SAMN56595522 |
| **sample01** | 1115 | *E. coli* | WGS | 30.6 | SAMN56595523 | - |
| **sample02** | 69 | *E. coli* | WGS | 30.6 | SAMN56595524 | - |
| **sample03** | 131 | *E. coli* | WGS | 30.6 | SAMN56595525 | - |
| **sample04** | 12 | *E. coli* | WGS | 30.6 | SAMN56595526 | - |
| **sample05** | 155 | *E. coli* | WGS | 30.6 | SAMN56595527 | - |
| **sample06** | 10 | *E. coli* | WGS | 30.6 | SAMN56595528 | - |
| **sample07** | 95 | *E. coli* | WGS | 30.6 | SAMN56595529 | - |
| **sample08** | 131 | *E. coli* | WGS | 30.6 | SAMN56595530 | - |
| **sample09** | 15061 | *E. coli* | WGS | 30.6 | SAMN56595531 | - |
| **sample10** | 69 | *E. coli* | WGS | 30.6 | SAMN56595532 | - |
| **sample11** | 69 | *E. coli* | WGS | 30.6 | SAMN56595533 | - |

**Table S4. Sequence types for routine clinical *E. coli*. The samples containing too few reads were excluded since they did not reach the threshold of 20X. Accession numbers for both phase 1 and phase 2 are displayed in the case of a DORAS experiments, otherwise only those for phase 1 (WGS) are displayed.**

| Sample | ST | Origin | Settings | Full resolution time (hours) | NCBI Accession number (Phase1, Phase2) | |
|---|---|---|---|---|---|---|
| **sample12** | 73 | *E. coli* | WGS | 30.6 | SAMN56595534 | - |
| **sample13** | 117 | *E. coli* | WGS | 30.6 | SAMN56595535 | - |
| **sample14** | 538 | *E. coli* | WGS | 30.6 | SAMN56595536 | - |
| **sample15** | 5640 | *E. coli* | WGS | 30.6 | SAMN56595537 | - |
| **sample16** | 567 | *E. coli* | WGS | 30.6 | SAMN56595538 | - |
| **sample17** | 69 | *E. coli* | WGS | 30.6 | SAMN56595539 | - |
| **sample18** | 131 | *E. coli* | WGS | 30.6 | SAMN56595540 | - |
| **sample19** | 131 | *E. coli* | WGS | 30.6 | SAMN56595541 | - |
| **sample20** | 1115 | *E. coli* | WGS | 30.6 | SAMN56595542 | - |
